# Introducing Rhythmic Sinusoidal Amplitude-Modulated Auditory Stimuli with Multiple Message Frequency Coding for Fatigue Reduction in Normal Subjects: An EEG Study

**DOI:** 10.1101/663344

**Authors:** Elham Shamsi, Zahra Shirzhiyan, Ahmadreza Keihani, Morteza Farahi, Amin Mahnam, Mohsen Reza Haidari, Amir Homayoun Jafari

## Abstract

Many of the brain-computer interface (BCI) systems depend on the user’s voluntary eye movements. However, voluntary eye movement is impaired in people with some neurological disorders. Since their auditory system is intact, auditory paradigms are getting more patronage from researchers. However, lack of appropriate signal-to-noise ratio in auditory BCI necessitates using long signal processing windows to achieve acceptable classification accuracy at the expense of losing information transfer rate. Because users eagerly listen to their interesting stimuli, the corresponding classification accuracy can be enhanced without lengthening of the signal processing windows. In this study, six sinusoidal amplitude-modulated auditory stimuli with multiple message frequency coding have been proposed to evaluate two hypotheses: 1) these novel stimuli provide high classification accuracies (greater than 70%), 2) the novel rhythmic stimuli set reduces the subjects’ fatigue compared to its simple counterpart. We recorded EEG from nineteen normal subjects (twelve female). Five-fold cross-validated naïve Bayes classifier classified EEG signals with respect to power spectral density at message frequencies, Pearson’s correlation coefficient between the responses and stimuli envelopes, canonical correlation coefficient between the responses and stimuli envelopes. Our results show that each stimuli set elicited highly discriminative responses according to all the features. Moreover, compared to the simple stimuli set, listening to the rhythmic stimuli set caused significantly lower subjects’ fatigue. Thus, it is worthwhile to test these novel stimuli in a BCI experiment to enhance the number of commands and reduce the subjects’ fatigue.

**Significance Statement:** Auditory BCI users eagerly listen to the stimuli they are interested in. Thus, response classification accuracy may be enhanced without the need for trial lengthening. Since humans enjoy listening to rhythmic sounds, this study was carried out for introducing novel rhythmic sinusoidal amplitude-modulated auditory stimuli with multiple message frequency coding. Our results show that each stimuli set evoked reliably discriminative responses according to all the features, and rhythmic stimuli set caused significantly lower fatigue in subjects. Thus, it is worthwhile to test these novel stimuli in a BCI study to increase the number of commands (by N^N^ permutations of just N message frequencies) and reduce the subjects’ fatigue.

## Introduction

Brain-computer interface (BCI) provides muscle-independent communication between brain and computer. Thus, translating the user’s intention to an external command (e.g., wheelchair control) comes true (Wolpaw et al., 2002). BCI can improve the quality of life for people with motor disabilities. Electroencephalogram (EEG) has been considered a reliable modality for using in BCI studies due to its noninvasiveness, good temporal resolution, easy implementation, and low cost (Wang et al., 2004; Hoffmann et al., 2008; Nijboer et al., 2008).

Patients with some neurological disorders, such as late-stage amyotrophic lateral sclerosis (ALS) and minimally conscious state (MCS), cannot perform voluntary eye movements or fixate their gaze. Moreover, daily usage of tactile BCI is hard because most people do not have tactile stimulators at home (Kaufmann et al., 2013). Thus, there has been an increasing interest towards auditory BCI (aBCI), which mainly uses auditory selective attention (Hill et al., 2004; Kanoh et al., 2008; Nijboer et al., 2008; Furdea et al., 2009; Klobassa et al., 2009; Kübler et al., 2009; Halder et al., 2010; Schreuder et al., 2010; Higashi et al., 2011; Höhne et al., 2011; Kim et al., 2011; Schreuder et al., 2011; Höhne et al., 2012; Kim et al., 2012; Lopez-Gordo et al., 2012b; Lopez-Gordo et al., 2012a; Käthner et al., 2013; Nakamura et al., 2013; Simon et al., 2014; Kleih et al., 2015; Halder et al., 2016; Zhou et al., 2016; Heo et al., 2017; Kaongoen and Jo, 2017; Jalilpour and Sardouie, 2018; Ogino et al., 2019) or auditory imagery (González et al., 2019) to influence event-related potentials (ERPs) and/or auditory steady-state responses (ASSRs). ASSR is chiefly evoked by listening to amplitude-modulated (AM) tones, and its spectrum has peaks at message frequency (f_m_) (Picton et al., 2003; Lopez et al., 2009; Tanaka et al., 2013; Tanaka et al., 2015).

In aBCI, lengthening the processing window enhances the classification accuracy, whereas it reduces the speed (Lopez-Gordo et al., 2012a). Also, when users eagerly listen to their interesting stimuli, the classification accuracy (Höhne et al., 2012; Treder et al., 2014; Zhou et al., 2016; Heo et al., 2017) and response amplitude (Kleih et al., 2010) rise. Moreover, rhythmic stimulation modulates the intrinsic neural oscillatory characteristics (Treder et al., 2014; Herrmann et al., 2016) and facilitates keeping focus (Sato et al., 2019). Interestingly, it was revealed that rhythmic (Keihani et al., 2018b; Keihani et al., 2018a) and chaotic (Shirzhiyan et al., 2019) visual alongside auditory stimuli (Shamsi et al., 2017) brought about both enough distinguishable EEGs and less subjects’ fatigue, compared to simple stimuli. Thus, rhythmic stimuli are one of the promising options for being used in aBCI. In some previous studies, single-message rhythmic SAM tones elicited EEG (Heo et al., 2017; Shamsi et al., 2017) and MEG (Kuriki et al., 2013). More specifically, response classification and subjects’ fatigue measurement were not performed in the study in which a sequence of AM tones was used to generate steady-state MEG (Kuriki et al., 2013). Further, responses were not classified in (Kuriki et al., 2013) and subjects’ fatigue were not evaluated in (Kuriki et al., 2013; Heo et al., 2017). Subjects’ preference and motivation play a major role in designing the experiments (Nijboer et al., 2008; Kleih et al., 2010; Ogino et al., 2019). Therefore, finding a way to reduce subjects’ fatigue without posing adverse effects on brain response discrimination is crucial. Unfortunately, shortening the stimuli duration might cause a decline in subjects’ fatigue at the expense of a reduction in response classification accuracy. However, innovative rhythmic stimuli are worth being designed and tested, thereby shedding more light on efficient fatigue reduction and response classification.

To our knowledge, there is not any research on AM sequences with multiple message frequencies. In this paper, six novel stimuli (i.e., simple and rhythmic sets) with multiple message frequency coding were introduced to test our hypotheses in healthy human subjects: 1) the resulting ASSRs are highly discriminative, and 2) listening to the novel rhythmic set reduces the subjective fatigue compared to the simple set.

## Materials and Methods

### Subjects

Nineteen healthy (twelve female) volunteers took part in this study. They all participated in our previous study (Shamsi et al., 2017), too. Their age was in the range of 22-29 years (25.26±2.05). All of them were right-handed according to Edinburgh Handedness Inventory (Oldfield, 1971) (Index: 0.75±0.26). Participants reported no musical expertise. The instructions were explained to them. Subjects signed written informed consent form before conducting the experiments. All the procedures were approved by the ethics committee and the deputy of research review board of Tehran University of Medical Sciences.

### Stimuli

In order to maintain consistency with other ASSR studies, double-sideband transmitted-carrier amplitude modulation with a modulation depth of 1 was used to generate the stimuli (to get more details, see (Lopez et al., 2009; Kuriki et al., 2013; Tanaka et al., 2013; Heo et al., 2017)):

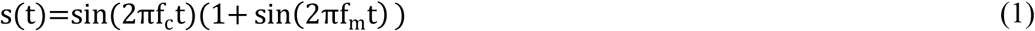

Where s(t) stands for the stimulus signal. In addition, f_c_ and f_m_ are carrier and modulation (i.e., message) frequency, respectively.

Two sets of stimuli were designed, each of which contained three stimuli. All the stimuli had a duration of 180 s. Each stimulus comprised three f_m_s. The stimuli sets are schematically represented in **Figure 1**. Using multiple message frequencies in each stimulus enriches its spectral content, because each f_m_ can elicit its corresponding peak in ASSR spectrum. In this way, different orders and permutations of f_m_s make it possible to produce various commands in aBCI. That is to say, using this kind of coding, only N message frequencies can generate N^N^ permutations, which means N^N^ stimuli and N^N^ commands, whereas N^N^ message frequencies in single-message sinusoidal AM tones are required for generating the same number of stimuli and commands. This is important because there is limitation for message frequency selection in the sense that strong ASSRs were elicited by message frequencies in the range of [30-50] Hz (Picton et al., 1987), so using multiple message frequency coding facilitates the construction of stimuli corresponding to possible commands.

**Figure 1.**
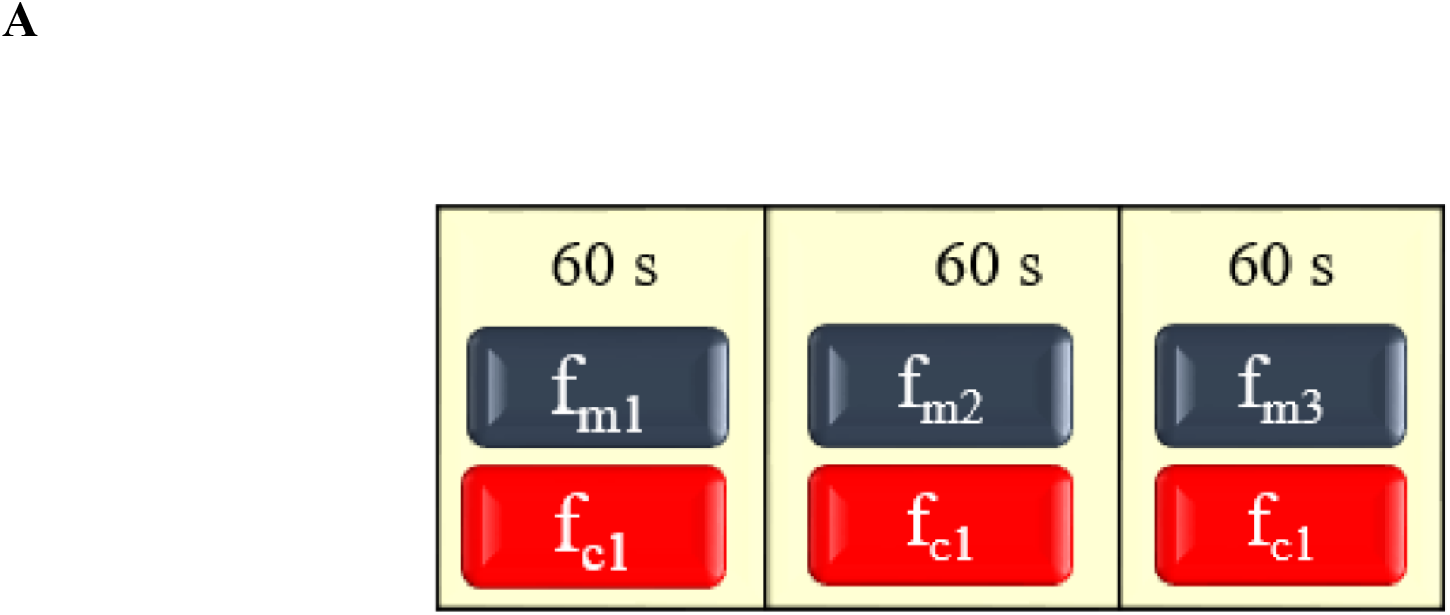
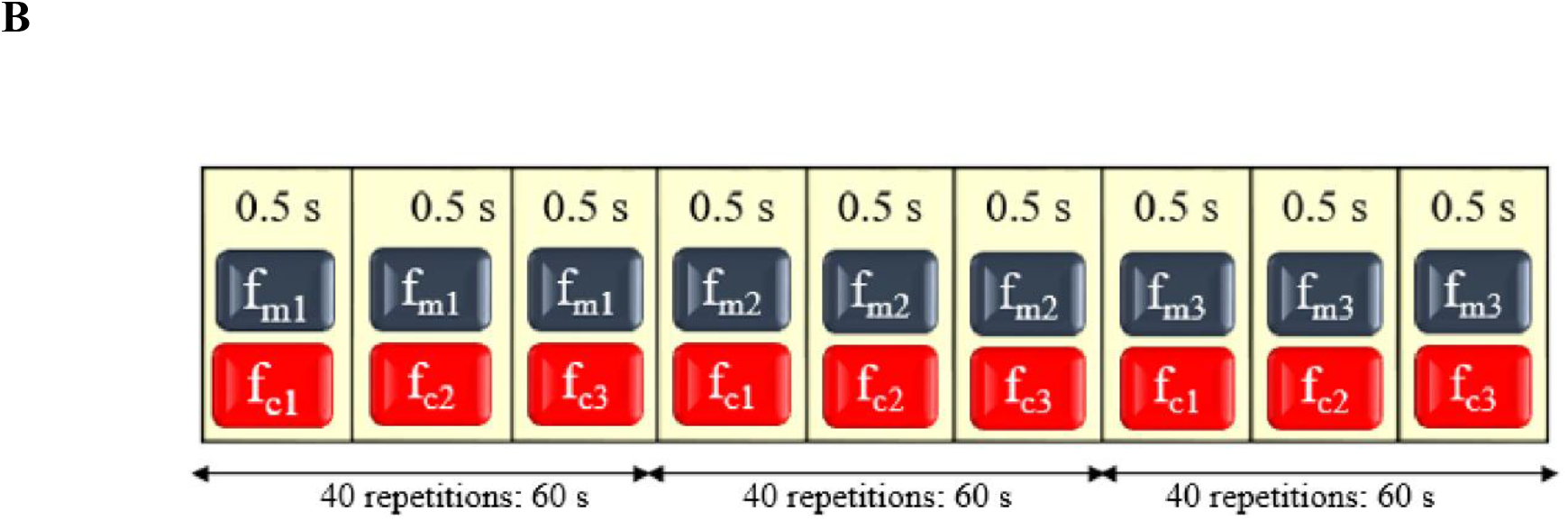
Schematic representation of the stimuli sets. ***A,*** simple set. ***B,*** rhythmic set.

It is noteworthy that in this paper, whenever only a single carrier was present in the stimuli, those stimuli are called *simple*, while the stimuli containing more than one carrier are referred to as *rhythmic*. In other words, *rhythm* was generated using multiple carriers. For both sets of stimuli, f_m_s were chosen to be among the (30, 35, 40) Hz. This is because consistent and robust ASSRs were elicited by message frequencies in the range of [30-50] Hz (Picton et al., 1987). Carrier frequencies were selected among the musical notes to be interesting for the subjects to listen to them. In this way, f_c_s were members of the (262, 392, 494) Hz corresponding to *do*, *sol*, and *si* musical notes, respectively. For the rhythmic stimuli, the presence of each carrier was set to 0.5 s according to the best tempo sensitivity time interval (Drake and Botte, 1993). Frequency details are displayed in **Table 1**. Each stimuli set contained ascending, descending, and one of the possible zigzagging codings of message/carrier frequency. In other words, within each set, we constructed only three permutations (out of 27 possible permutations).

**Table 1.** Message and carrier frequency details in the stimuli sets

|  |  | Frequency (Hz) |  |  |  |  |  |
| --- | --- | --- | --- | --- | --- | --- | --- |
| | | $f_{m1}$ | $f_{m2}$ | $f_{m3}$ | $f_{c1}$ | $f_{c2}$ | $f_{c3}$ |
| Stimuli name |  |  |  |  |  |  |  |
| Simple set | SA | 30 | 35 | 40 | 262 | - | - |
|  | SB | 40 | 35 | 30 | 494 | - | - |
|  | SC | 40 | 30 | 35 | 392 | - | - |
| Rhythmic set | RA | 30 | 35 | 40 | 494 | 262 | 392 |
|  | RB | 40 | 35 | 30 | 262 | 392 | 494 |
|  | RC | 40 | 30 | 35 | 494 | 392 | 262 |

For instance, a pattern with 30-30-35 Hz (two identical f_m_s at first and second portions of the triple pattern) or any similar permutation was not constructed. The reason is that we wanted to see the distinguishability that all of 30, 35 and 40 Hz within the proposed coding can provide in the corresponding ASSRs. Therefore, the presence of all three f_m_s in the proposed coding was required in this study. All the stimuli were generated in MATLAB R2016b (MathWorks Inc., Natick, MA, USA). Sampling frequency for all the stimuli was 4410 Hz.

### Task

In order to assess the subjects’ level of some psychological factors (i.e., depression, stress, anxiety) through the week before the experiment, they were asked to fill 42-item depression, anxiety, and stress scales (DASS-42) questionnaire (Lovibond and Lovibond, 1995) preceding the stimuli presentation. Also, the subjects filled questionnaire of current motivation (QCM) (Rheinberg et al., 2001; Vollmeyer and Rheinberg, 2006) before the beginning of the experiment. In this way, it was possible to measure their motivation and interest for participation, sense of challenge about the task, and anxiety they feel about the task. Then, the participants were requested to remain eyes-closed, still, and listen to the stimuli. For preventing from effects of the stimuli presentation order on the reported fatigue, the stimuli were presented in a random order for each participant. In other words, stimuli presentation order differed between the subjects. After listening to each stimulus, participants reported the amount of stimulus-induced fatigue that they experienced, as an integer number from 0 (minimum fatigue) to 10 (maximum fatigue) according to the visual analog scale (VAS) and they were given a short break of 60-120 seconds before presentation of the next stimulus. This procedure was performed for every stimulus. There were two reasons for this separate presentation: 1) we wanted to ensure that the stimuli in each set (i.e., simple, rhythmic) elicit sufficient inherently distinguishable responses in the brain, 2) we wanted to measure the amount of fatigue that each stimulus caused to each subject, so we had to present the stimuli separately (i.e., one by one). The stimuli presentation procedure is displayed in **Figure 2**. It is worth mentioning that all the previously mentioned psychological data (i.e., obtained via DASS-42 and QCM questionnaires) were later used to explore whether there is a relationship between those factors and the fatigue level that subjects reported. In other words, these data helped us validate whether the reported fatigue was solely related to the stimuli.

**Figure 2.**
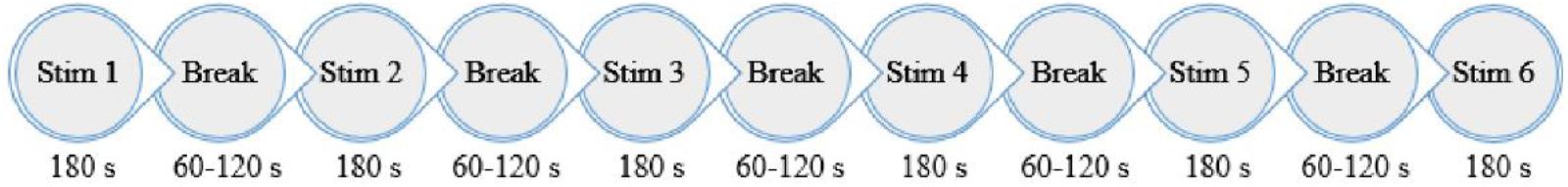
Stimuli presentation procedure. *Stim* is the abbreviation of stimulus. The stimuli were presented in a random order. After listening to each stimulus, which was 180 s long, the subjects reported the level of stimulus-induced fatigue that they experienced, as an integer number from 0 (minimum fatigue) to 10 (maximum fatigue) according to VAS. Then, they were given a short break of 60-120 s before presentation of the next stimulus.

### Experiment apparatus and recording

Insert earphones ER-3A (Etymotic Research, Elk Grove Village, IL) presented the stimuli to the subjects. For each stimulus, the volume was set according to equal loudness level contours at the standard ISO 226:2003.

Electrode placement was performed according to 10-20 international system. Active g.LADYbird electrodes were placed on Fz, Cz, T7 and T8. The reason for selecting Fz was that it is shown that Fz has the highest average amplitude of responses in subjects to whom a music is pleasant (Kayashima et al., 2017). Three other channels were consistently used in a number of previous ASSR studies (Lopez et al., 2009; Higashi et al., 2011; Kim et al., 2011; Heo et al., 2017; Shamsi et al., 2017). Right earlobe and Fpz were considered as the reference and ground, respectively (Heo et al., 2017; Shamsi et al., 2017). EEG was recorded by g.USBamp (g.tec Medical Engineering GmbH, Austria) at a sampling frequency of 4800 Hz. Online filters consisted of a bandpass with a bandwidth of [0.5-2000] Hz, and a notch with a center frequency of 50 Hz.

### Signal analysis

Firstly, in order to detect ASSR, we explored whether the amplitude spectrum at f_m_ was larger than the mean+3×standard deviation (SD) of the amplitude spectrum at frequencies in the range of (f_m_-1 to f_m_-5) and (f_m_+1 to f_m_+5) (Tanaka et al., 2015). Then, prominent features (i.e., power spectral density (PSD), Pearson’s correlation coefficient (PCC), and canonical correlation coefficient (CCC)) were extracted to be fed into the classifier. All the analyses were carried out in MATLAB R2016b (MathWorks Inc., Natick, MA, USA) on a laptop, which had Intel^®^ Core™ i7-2670QM CPU @ 2.20 GHz as its processor.

### Power spectral density

Keeping in mind that PSD is a robust feature for analyzing ASSR, it was computed (using amplitude spectrum at f_m_ and its adjacent frequencies in the range of f_m_−5-f_m_−1 and f_m_+1-f_m_+5) according to the literature (Tanaka et al., 2013; Tanaka et al., 2015), as follows:

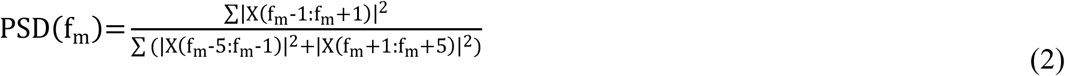

Where |X(f_m_)| is the amplitude spectrum of the brain response at frequency of f_m_.

### Pearson’s correlation coefficient

To investigate the amount of correlation between each stimulus and its corresponding ASSR, Pearson’s correlation coefficient was used, which, through this paper, will be referred to as *PCC*. As previously mentioned, the spectrum of this response has a peak at the modulation frequency (f_m_) (Tanaka et al., 2013; Tanaka et al., 2015), which is exactly the same as the fundamental frequency of the stimulus envelope. Thus, PCC was calculated for investigating the amount of correlation between each stimulus envelope (i.e., X) and the ASSR (i.e., Y) that the stimulus elicited. It was calculated as follows:

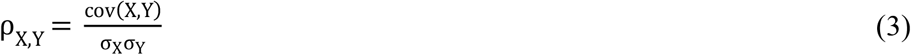

Where cov(X,Y), σ_X_, and σ_Y_ are the covariance of the two signals, the standard deviation of X and the standard deviation of Y, respectively. ρ_X,Y_ has a value within the interval of [−1, 1]. Obviously, if there is not any linear relationship between X and Y, it will be zero.

### Canonical correlation coefficient

This method seeks for a pair of linear combinations for two signals in such a way that the correlation between the two canonical signals becomes maximized. In this way, pairs having linear combinations with the most linear correlation are chosen in a way that the previously identified pairs are orthogonal to them. If the EEG is represented by X and the stimulus envelope is considered to be Y, their projection vectors are denoted by 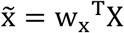 and 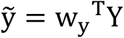, respectively. Solving the equation below, w_x_ and w_y_ can be obtained:

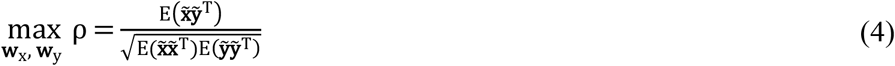

Where, ρ is called the canonical correlation coefficient.

### Classification

There were two cases for the classification, one for the responses to simple stimuli set, and the other for those of the rhythmic stimuli set. That is to say, in this study, two three-class response classification existed. Each of these two classifications was performed three times, based on whether PSD, PCC, or CCC was fed into the classifier. Feature extraction and classification were performed across every 20-s segment of the EEGs to be consistent with the relevant literature and practical applications (Kim et al., 2011; Heo et al., 2017). Therefore, for each subject and their responses to each stimuli set, there were 4 channels × 9 segments × 3 classes = 108 observations. Classification was conducted by means of five-fold cross-validated naïve Bayes classifier. The chosen classifier utilizes the total probability theorem and the Bayes theorem to estimate the posterior probability (i.e., the probability that the features of an observation belong to a particular class) for each class. Then, for each observation, corresponding posterior probabilities are compared to each other and the highest will be selected as the outcome of the classification. Naïve Bayes classifier performs classification on the assumption that features of each class have statistical independence, whereas sometimes this is not the case. This classifier, however, works well in practice (Hastie et al., 2009). Posterior probability was calculated as follows:

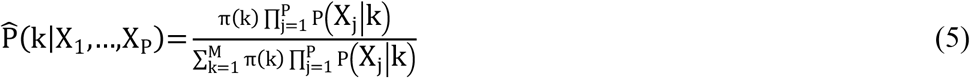

Where, k is the class index, X_1_,…,X_P_ are the features for each observation, and π(k) is the empirical prior probability of class k. It is worth mentioning that the hyperparameters of the naïve Bayes classifier for each training fold was determined by Bayesian optimization.

In order to evaluate the amount of classification performance, classification accuracy and Cohen’s kappa value were computed. Classification accuracy was defined to be the number of correctly classified observations divided by the number of classified observations. According to (Billinger et al., 2012), Cohen’s kappa value was calculated as follows:

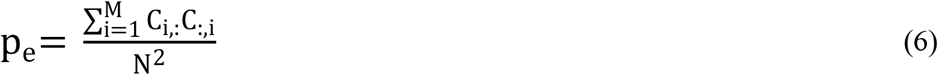

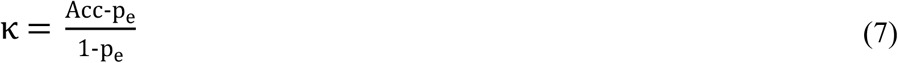

Where, p_e_ is the chance level, C_i,:_ is the i-th row of confusion matrix, C_:,i_ is the i-th column of confusion matrix, M represents the number of classes, and N is the total number of classified observations. Besides, κ and Acc are the Cohen’s kappa value and the classification accuracy, respectively.

### Experimental Design and Statistical Analysis

In order to examine the effect of stimuli type (i.e., simple, rhythmic) and feature (i.e., PSD, PCC, CCC) on the classification performances, we used the classification performance measures (i.e., accuracy, Cohen’s kappa value) for all the subjects (12 female, 7 male) as the dependent variable. Features and stimuli types were the within-subjects factors. All these were carried out via within-subjects repeated measures ANOVA. Mauchly’s test checked whether the sphericity assumption was held. Further, Greenhouse-Geisser approximation corrected the degrees of freedom. We selected Tukey’s honest significant difference to perform *post hoc* comparisons.

In addition, Wilcoxon signed rank test was used to see whether the level of stimuli-induced fatigue corresponding to the two sets of stimuli differed significantly within the subjects. In this statistical design, subjects’ fatigue was the dependent variable and type of the stimuli sets (i.e., simple and rhythmic) were within-subjects factors.

To investigate whether there is a relationship between psychological factors, which were evaluated via the questionnaires, and the reported fatigue, Spearman’s correlation test was performed. All the analyses were conducted in MATLAB R2016b (MathWorks Inc., Natick, MA, USA). Signal recording and analyses procedure is illustrated in **Figure 3**.

**Figure 3.**
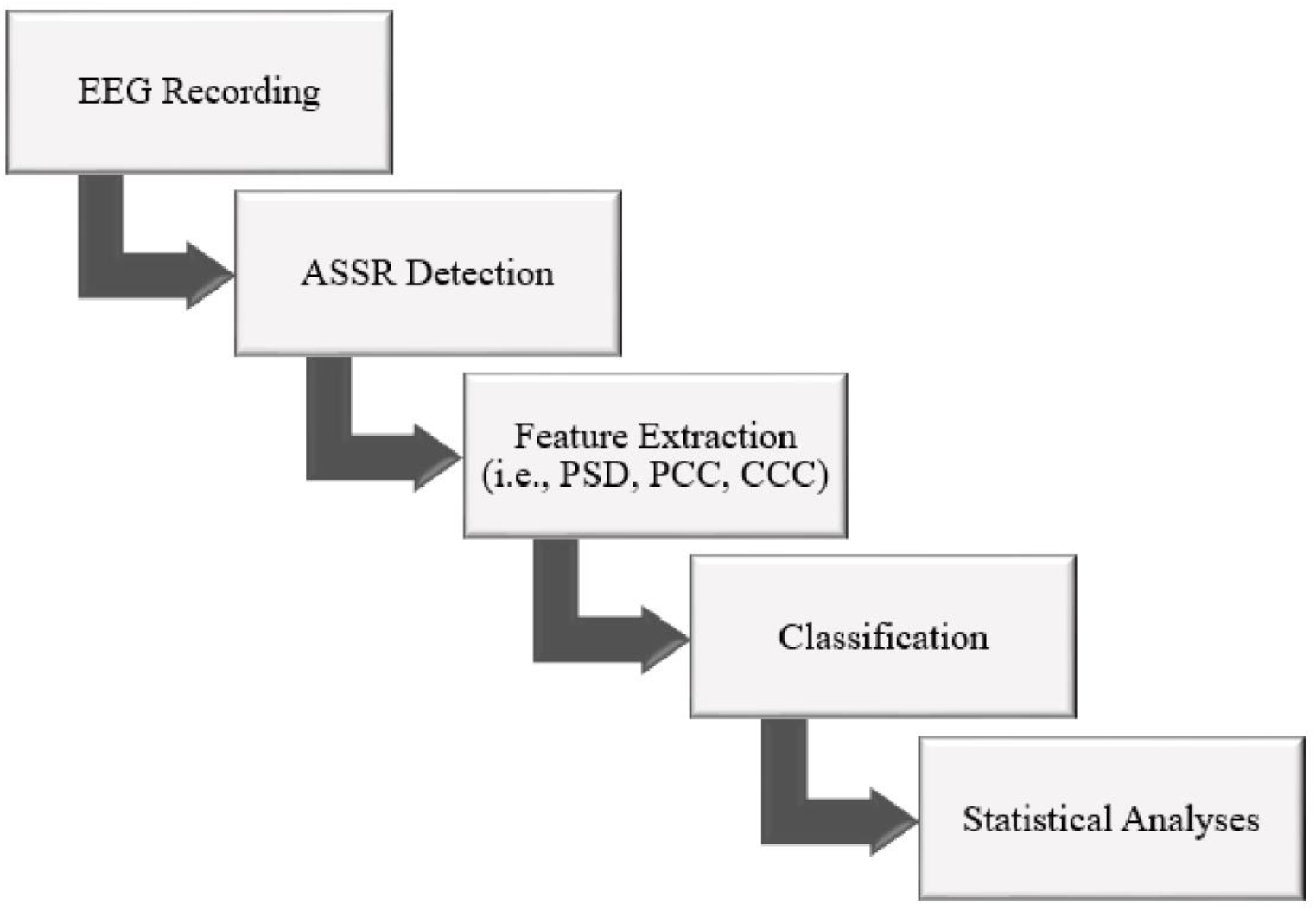
Signal recording and analyses procedure.

## Results

### Response detection

For each stimulus, amplitude spectrum of its corresponding EEG was computed. We checked whether the amplitude spectrum at f_m_ was larger than the mean;3×standard deviation (SD) of the amplitude spectrum at frequencies in the range of (f_m_−1 to f_m_−5) and (f_m_+1 to f_m_+5) (Tanaka et al., 2015) to make sure that ASSR was appeared. For simple and rhythmic stimuli set, amplitude spectrum corresponding to one stimulus is denoted in **Figure 4.** as a representative, because all the ASSRs satisfied the aforementioned condition (Tanaka et al., 2015). Consequently, these newly designed auditory stimuli with multiple message frequency coding elicited robust ASSRs.

**Figure 4.**
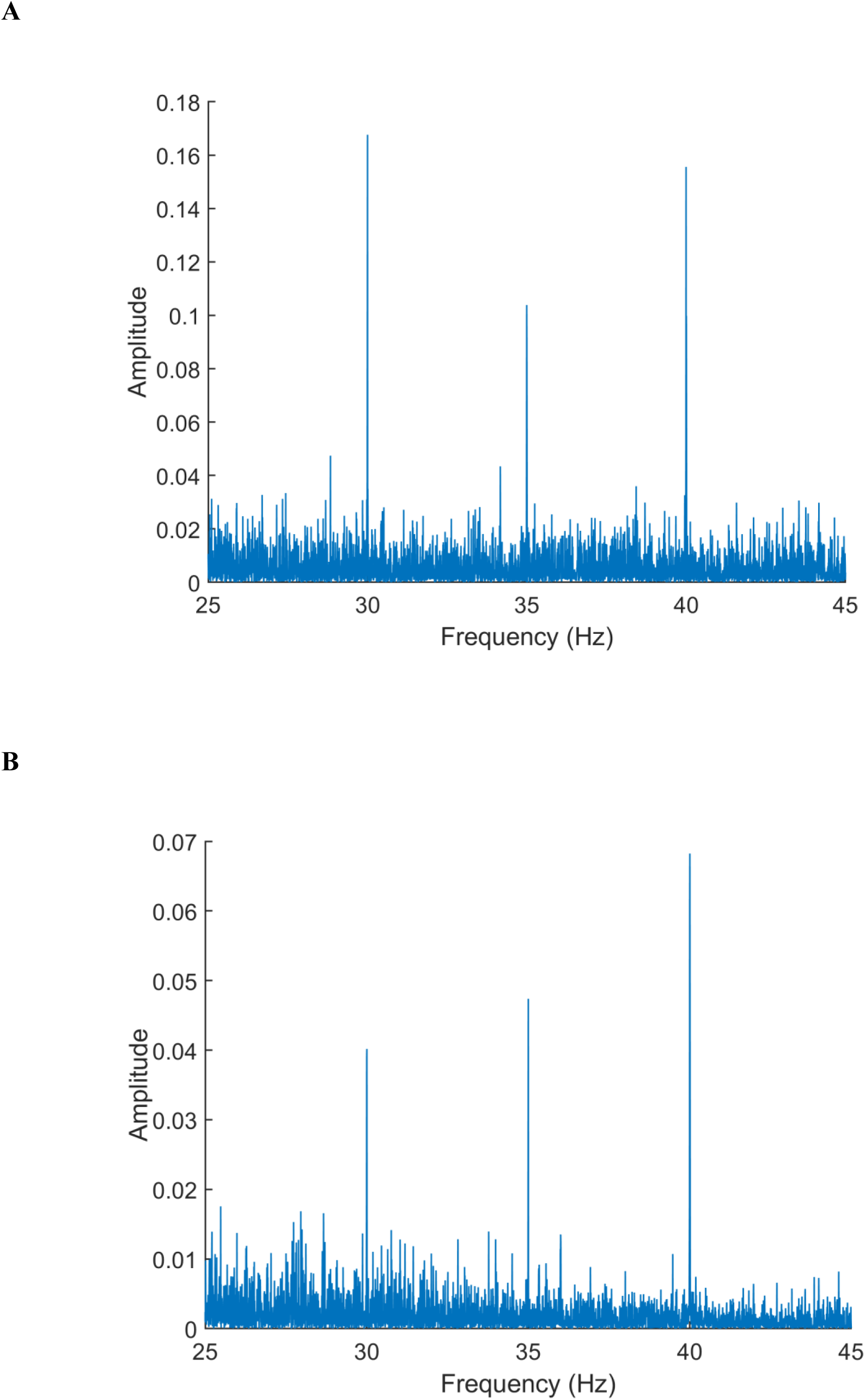
Amplitude spectrum of the responses to representative stimuli ***A,*** simple (SA). ***B,*** rhythmic (RA).

### Behavioral results

After listening to each stimulus, participants reported the amount of subjective fatigue that they experienced by listening to that stimulus. All the subjects reported their level of fatigue as an integer number in the range from 0 (minimum) to 10 (maximum) according to VAS. In comparison to the simple stimuli, the rhythmic stimuli caused significantly lower fatigue (*p* = 0.005, Wilcoxon signed rank test). This confirms our second hypothesis (fatigue reduction using the proposed novel rhythmic stimuli set). Boxplot representation of fatigue due to the stimuli in each set is illustrated in **Figure 5**.

**Figure 5.**
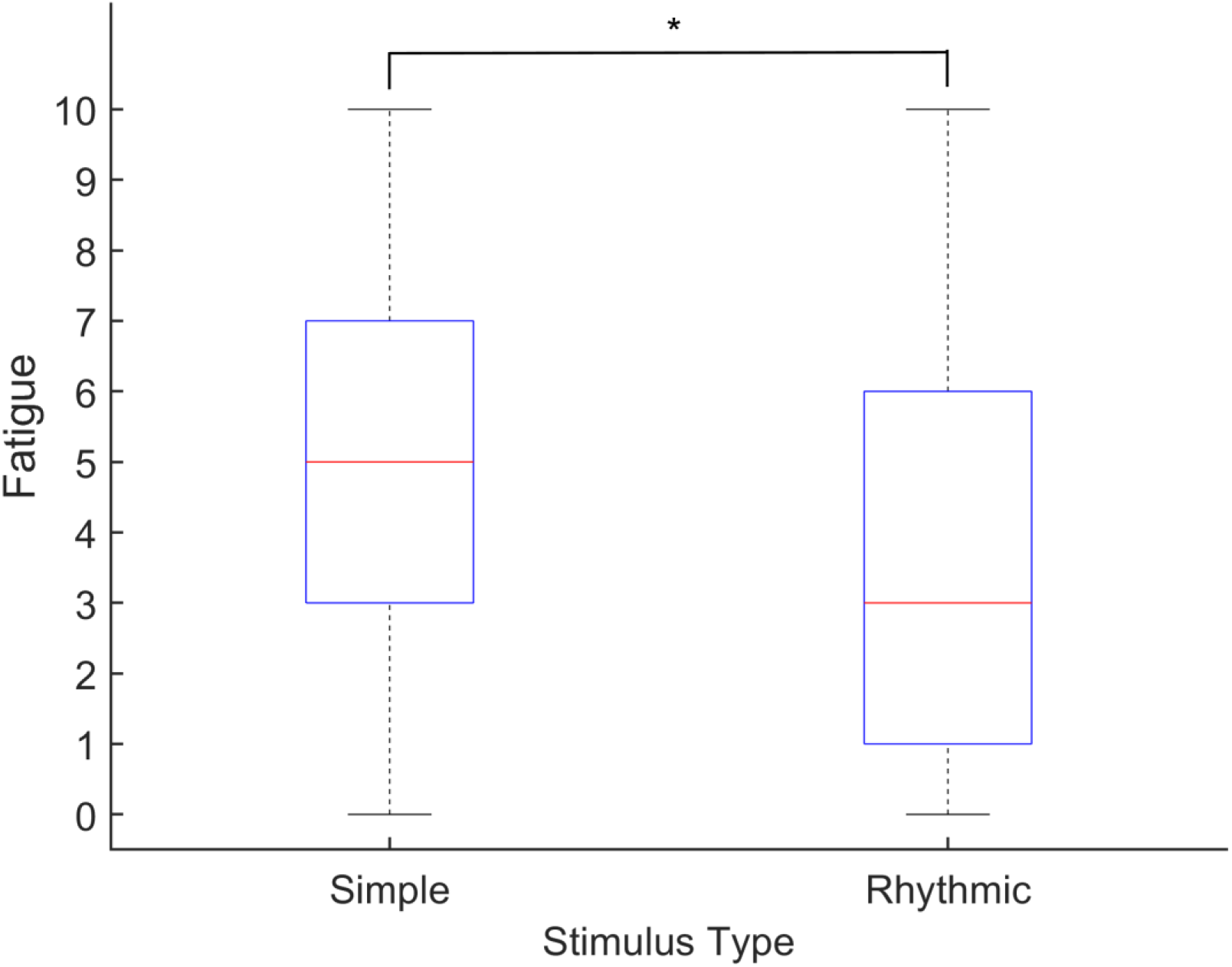
Boxplot representation of fatigue caused by the stimuli (fatigue levels were integers from 0 (minimum fatigue) to 10 (maximum fatigue) according to VAS). Rhythmic set significantly reduced the subjects’ fatigue (*p* = 0.005, Wilcoxon signed rank test).

In addition, there was no significant correlation between psychological factors and fatigue caused by listening to each stimulus. This is an indication of the fact that the subjects truly reported the fatigues that were mainly caused by listening to the stimuli, regardless of their psychological state. For each stimulus, scatter plot of the data that yielded strongest Spearman’s correlation coefficient, along with the best monotone curve fitted to the data are denoted on **Figure 6.** In each case, the best-fit curve was obtained through nonlinear least squares method. Details of all the Spearman tests (Spearman’s correlation coefficients and corresponding *p*-values) are illustrated **Table 2**.

**Table 2.** Spearman’s correlation test (r: Spearman’s correlation coefficient, P: *p* obtained via the test). There was not any significant correlation between the psychological factors and stimuli-caused fatigue

|  | Challenge | Interest | Success<br>Probability | Anxiety<br>(QCM) | Depression | Anxiety<br>(DASS) | Stress |
| --- | --- | --- | --- | --- | --- | --- | --- |
| Fatigue Caused by SA | r = -0.088<br>P = 0.721 | r = -0.055<br>P = 0.823 | r = 0.099<br>P = 0.685 | r = 0.086<br>P = 0.726 | r = -0.122<br>P = 0.619 | r = -0.098<br>P = 0.691 | r = -0.217<br>P = 0.372 |
| Fatigue Caused by SB | r = 0.049<br>P = 0.843 | r = 0.185<br>P = 0.449 | r = 0.086<br>P = 0.726 | r = 0.114<br>P = 0.643 | r = 0.025<br>P = 0.920 | r = 0.263<br>P = 0.276 | r = 0.015<br>P = 0.952 |
| Fatigue Caused by SC | r = 0.149<br>P = 0.543 | r = 0.314<br>P = 0.191 | r = 0.109<br>P = 0.658 | r = 0.084<br>P = 0.736 | r = -0.127<br>P = 0.603 | r = 0.176<br>P = 0.470 | r = -0.101<br>P = 0.680 |
| Fatigue Caused by RA | r = -0.074<br>P = 0.765 | r = -0.160<br>P = 0.512 | r = -0.055<br>P = 0.823 | r = 0.015<br>P = 0.951 | r = 0.071<br>P = 0.772 | r = 0.268<br>P = 0.267 | r = -0.006<br>P = 0.981 |
| Fatigue Caused by RB | r = -0.012<br>P = 0.962 | r = -0.190<br>P = 0.435 | r = -0.047<br>P = 0.848 | r = 0.170<br>P = 0.488 | r = 0.371<br>P = 0.118 | r = 0.454<br>P = 0.051 | r = 0.308<br>P = 0.200 |
| Fatigue Caused by RC | r = 0.176<br>P = 0.472 | r = -0.070<br>P = 0.776 | r = -0.298<br>P = 0.216 | r = 0.049<br>P = 0.844 | r = 0.247<br>P = 0.308 | r = 0.438<br>P = 0.061 | r = 0.413<br>P = 0.079 |

**Figure 6.**
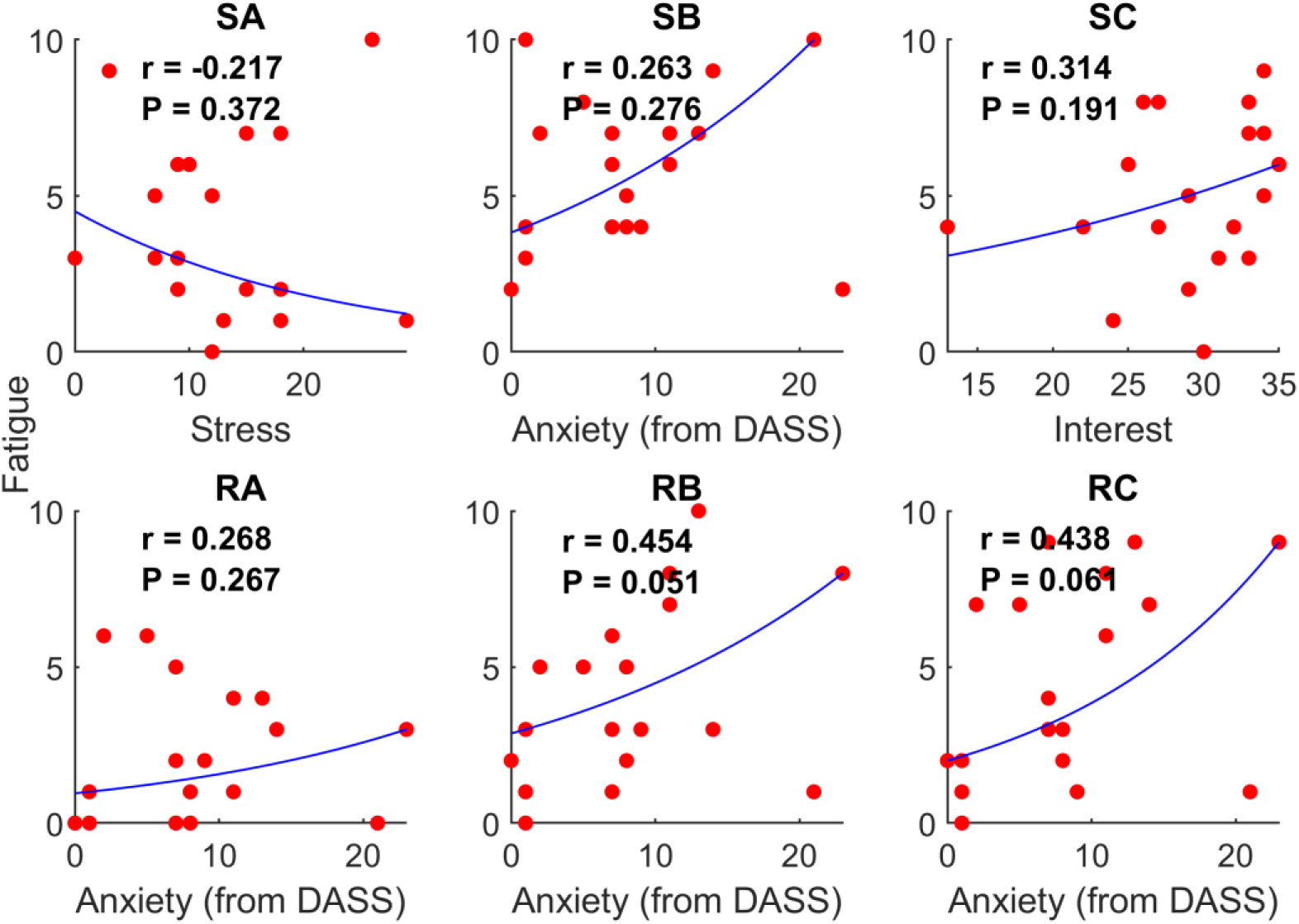
Scatter plots of the data that yielded strongest Spearman’s correlation coefficient between the psychological factors and fatigue caused by each stimulus (each plot’s name), along with the best monotone curve fitted to the data. There was not any significant correlation between psychological factors and fatigue caused by listening to each stimulus. In each plot, r represents Spearman’s correlation coefficient, while P is the *p* obtained in the Spearman’s correlation test.

### Classification performance

To investigate whether the designed stimuli evoke intrinsically discriminative responses, we performed response classification (for details on the features and classifier parameters used, see Materials and Methods). For both sets of stimuli, high classification accuracy and Cohen’s kappa value, up to a maximum of 100% and 1, respectively, were obtained. There was no significant difference between the responses to the simple and rhythmic stimuli sets in terms of classification performance (F(1,15) = 4.06, *p* = 0.062, repeated measures ANOVA). Furthermore, there was no significant difference between PSD, PCC, and CCC features in terms of classification performance (F(2,30) = 1.21, *p* = 0.307, repeated measures ANOVA). These indicate that the responses to the stimuli in each set are sufficiently discriminative. Further, the results show that all the extracted features were discriminant measures for the responses to each set. Group means of the classification accuracy and Cohen’s kappa value for different conditions (i.e., features) as a function of the stimuli type (i.e., simple, rhythmic) are illustrated in **Figure 7**. The results show that all the average classification accuracies are well above 70%, which is the minimum acceptable classification accuracy in BCI systems. Thus, the responses to the stimuli in each set are highly discriminative. In other words, our first hypothesis is confirmed. To investigate whether there is generalizability in terms of response discrimination, we also performed between-subjects classification for the responses to each set. Classification accuracy and Cohen’s kappa value for between-subjects classification according to each feature are listed in **Table 3**.

**Table 3.** Between-subjects classification performance for different stimuli sets and features

|  | PSD |  | PCC |  | CCC |  |
| --- | --- | --- | --- | --- | --- | --- |
|  | Simple | Rhythmic | Simple | Rhythmic | Simple | Rhythmic |
| Accuracy (%) | 92.98 | 88.89 | 93.57 | 81.29 | 94.15 | 83.04 |
| Cohen's kappa value | 0.90 | 0.83 | 0.90 | 0.72 | 0.91 | 0.75 |

**Figure 7.**
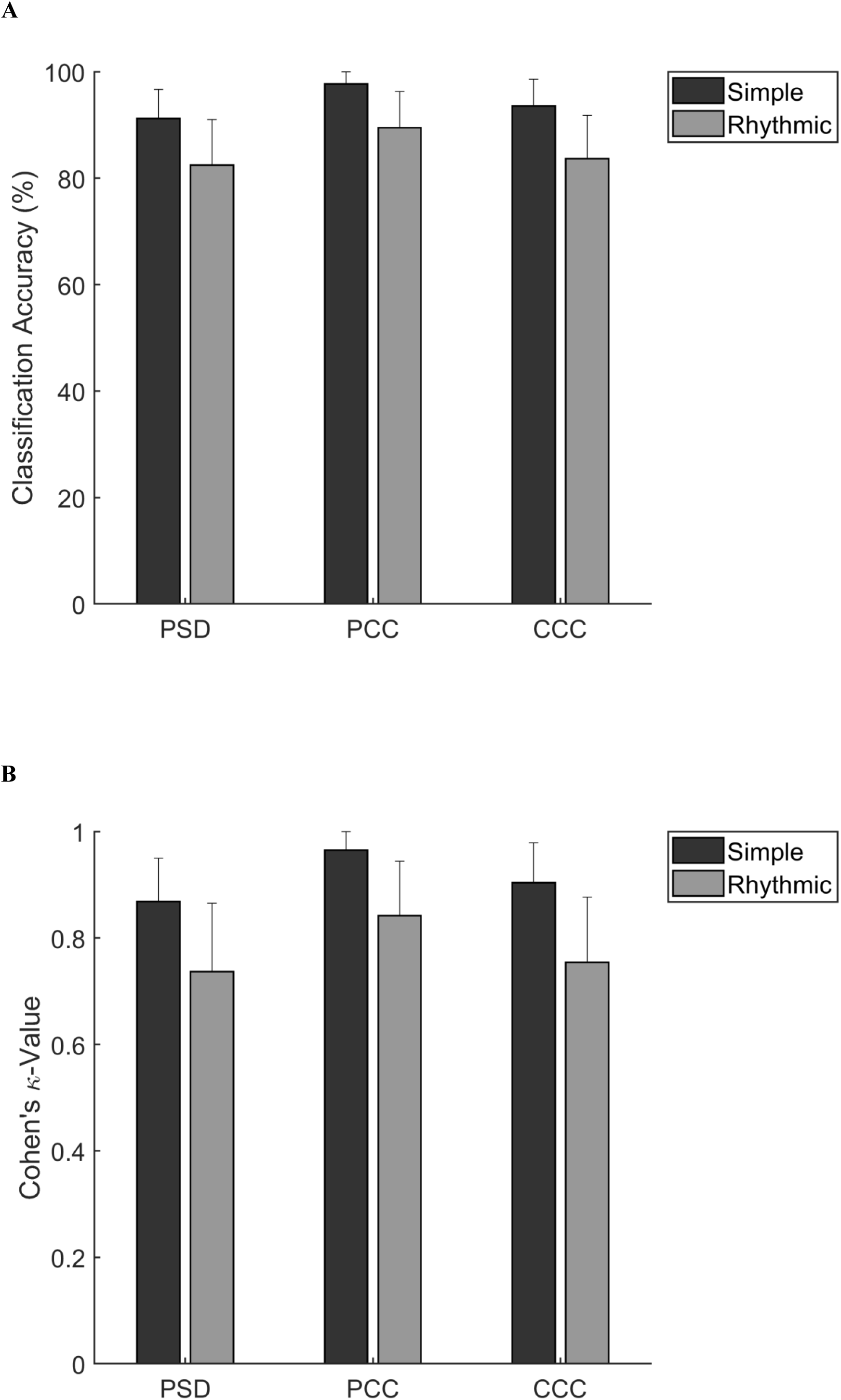
***A,*** Group means of the classification accuracy for the three features (PSD, PCC, CCC) as a function of stimuli type (illustrated as Simple, Rhythmic). ***B,*** Group means of the Cohen’s kappa value for the three features (PSD, PCC, CCC) as a function of stimuli type (illustrated as Simple, Rhythmic).

## Discussion

Finding a way to reduce subjects’ fatigue without posing adverse effects on brain response discrimination is crucial. As humans enjoy listening to rhythmic sounds, it seems that utilizing rhythmic auditory stimuli in the experiments that aim to evoke and examine auditory responses in the brain might reduce the subjects’ fatigue. Additionally, sinusoidal amplitude-modulated tones are helpful in studies that have encoding of envelope and periodicity in human’s auditory system. Moreover, they can also be used in ASSR-based BCI systems. Therefore, exploring SAM tone-evoked ASSR is important. This study was carried out to test our two hypotheses in healthy humans: 1) the ASSRs to the novel stimuli with multiple message frequency coding are highly discriminative, and 2) listening to the novel rhythmic stimuli set with multiple message frequency coding reduces the subjective fatigue. All these were conducted to determine whether the stimuli introduced in this paper have enough feasibility, in terms of classification performance and subjective fatigue, to be used in an aBCI. Both of the stimuli sets designed in this study were novel in the sense of having multiple message frequency coding. In addition, response classification and fatigue evaluation were carried out for these stimuli for the very first time. For a better insight into the novelty of this work, a comparison between the results of this paper and those of some relevant ASSR studies that performed response classification, are illustrated in **Table 4**.

**Table 4.**
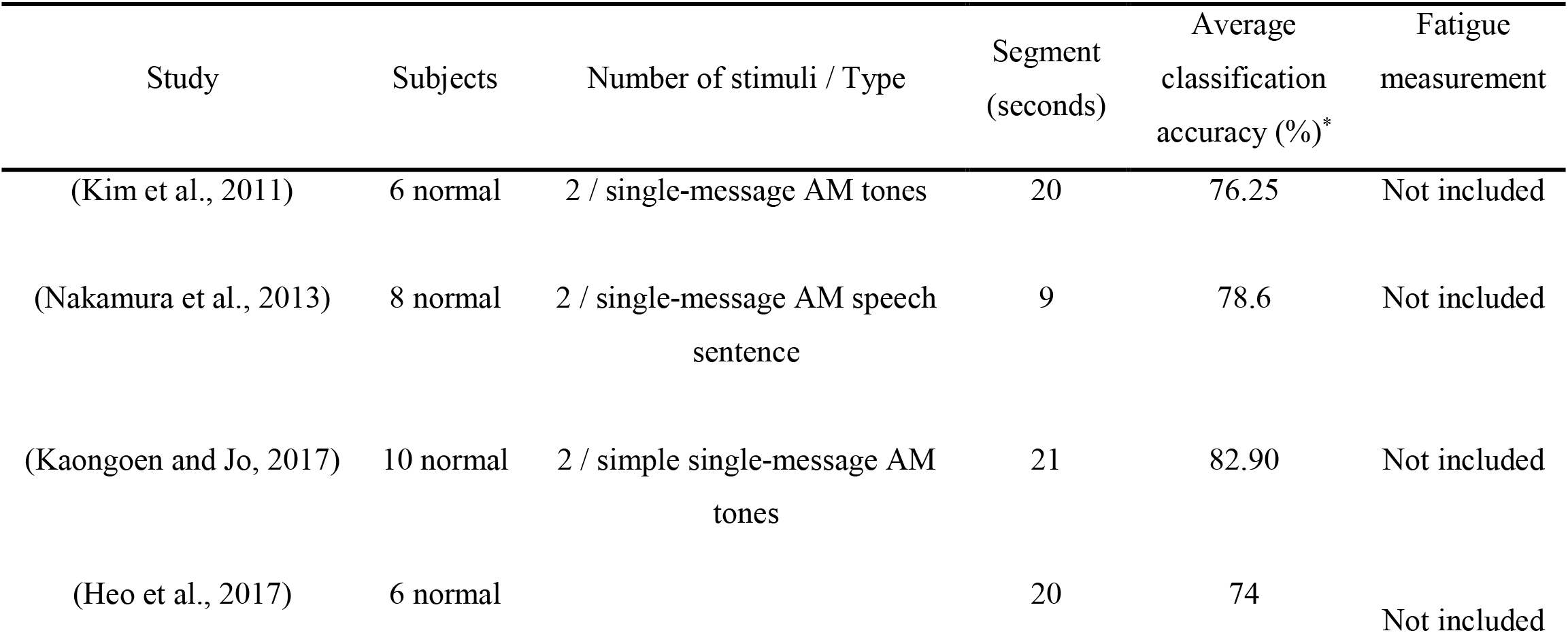
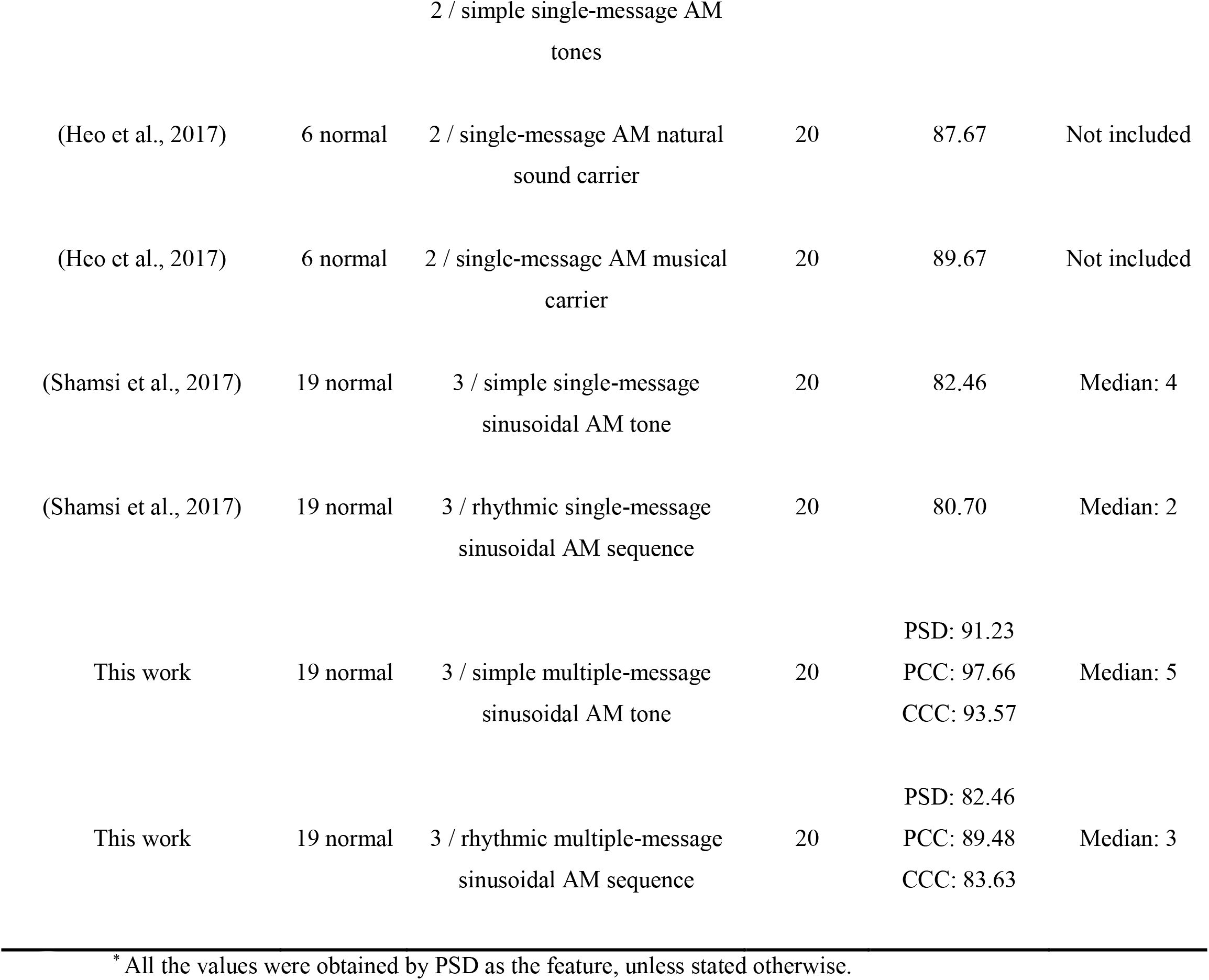
A comparison between the results of this paper and those of some relevant ASSR studies

Robust peak in ASSR spectrum at message frequencies (i.e., corresponding to the envelope of the stimuli) is consistent with previous findings (Lopez et al., 2009; Kim et al., 2011; Nozaradan et al., 2012; Kuriki et al., 2013; Kaongoen and Jo, 2017). It confirms the notion that human auditory system acts like an envelope detector, which can be an amplitude demodulator (Miyazaki et al., 2013). In addition, amplitudes of the responses to the rhythmic set were lower than those of the simple set. This result, at first glance, seems to contradict the findings of another study showing that interesting stimuli lead to enhanced response amplitudes (Kleih et al., 2010). However, our study measured the subjects’ fatigue rather than their acceptance rate or feeling about the stimuli. Therefore, it is not perfectly clear whether the subjects liked the rhythmic set. Lower response amplitude obtained by rhythmic stimuli set might occurred due to the more complex structure of the rhythmic set, compared to that of the simple set. There is relevant supporting evidence that the more complex stimuli elicited less amplitude, when compared to stimuli with simpler structure (Nakamura et al., 2013; Shamsi et al., 2017).

Moreover, the rhythmic stimuli set resulted in significantly less fatigue in the subjects, compared to that of the simple stimuli set. This is in agreement with the findings of our previous study on the comparison between the fatigue levels that simple and rhythmic single-message sinusoidal AM stimuli can cause (Shamsi et al., 2017). It is also in accordance with the results of previous studies on rhythmic (Keihani et al., 2018b; Keihani et al., 2018a) and chaotic (Shirzhiyan et al., 2019) visual stimuli that reduced the subjective fatigue, and confirms our second hypothesis. In addition, the insignificant and infinitesimal correlation between the fatigue and the psychological state can ensure us that the subjects truly reported the fatigue that was mainly caused by listening to the stimuli, regardless of their psychological status.

We were able to perform highly accurate, precise and reliable classification on within- and between-subjects responses without any artifact rejection. This shows that there was adequate inherent discrimination even at the raw signal level for the responses to each stimuli set. It can be seen from the within-subjects classification performance results that firstly, stimuli with multiple message frequencies generate highly distinguishing ASSRs. They, thus, have the potential to be utilized in aBCI to increase the number of available commands, and therefore the information transfer rate. This can take place by means of multiple permutations of just a few message frequencies, which is accessible via the coding presented in this paper. In other words, fewer message frequencies (i.e., N in the proposed multiple message coding, compared to N^N^ in single-message SAM tones) can generate N^N^ commands in aBCI. Secondly, the rhythmic stimuli elicit discriminative responses, which are as distinct as that of the simple stimuli. Thirdly, all the extracted features in this study (i.e., PSD, PCC, and CCC) are discriminant measures for the classification of the ASSR to the stimuli with multiple message frequencies. Additionally, high amounts of between-subjects classification performance indicate that the ASSRs to the stimuli in each set were reliably distinct and generalizable. Furthermore, all the average classification accuracies were far above 70%, which is sufficient for a BCI system. In other words, our first hypothesis was also confirmed. Thus, the stimuli designed in this paper have the adequate potential to be corresponding to several different commands and to generate distinct responses in BCI systems.

Although both stimuli sets resulted in sufficient classification performances, rhythmic set led to insignificant lower classification accuracy and Cohen’s kappa value than its simple counterpart. This result, at first glance, seems to be opposite to the findings in the literature that interesting stimuli provide higher response classification accuracy (Höhne et al., 2012; Treder et al., 2014; Zhou et al., 2016; Heo et al., 2017). We measured the subjects’ fatigue rather than their acceptance rate or feeling about the stimuli. Consequently, it is not perfectly clear whether the subjects liked the rhythmic set. Insignificant lower classification performance achieved by rhythmic stimuli set might have taken place due to the more complex structure of the rhythmic set. The fact that the more complex stimuli elicited less amplitude, when compared to stimuli with simpler structure (Nakamura et al., 2013; Shamsi et al., 2017) might also have led to lower classification performance.

The average classification performances obtained in this study outperformed previous studies, which utilized single-message AM tones (Lopez et al., 2009; Kim et al., 2011; Heo et al., 2017; Kaongoen and Jo, 2017; Shamsi et al., 2017) and single-message AM sentences (Nakamura et al., 2013). Particularly, the average classification performances obtained for our simple set was higher than those of a research, which used single-message AM natural sound carriers (Heo et al., 2017). However, the average classification performances for our rhythmic set was a bit lower than those of the same study, which also made use of single-message AM instrumental music carriers (Heo et al., 2017). These differences may relate to the different experimental protocols, including stimulus characteristics and sample size (**Table 4**).

The results showed that stimuli in each set have sufficient inherent discrimination to the extent that it is worthwhile to use these novel auditory stimuli with multiple message frequency coding in a BCI experiment. If we are requested to choose between our proposed stimuli sets to be utilized in BCI studies, the choice will be the rhythmic set. The reason is listening to the rhythmic set reduced the subjects’ fatigue and the brain responses to the rhythmic set were classified via a common classifier, with nearly as high performance as the simple set. This set will also be able to increase the number of possible commands by permutation of the message frequencies within its stimuli.

## Limitations and Future Directions

In this paper, each stimuli set contained ascending, descending, and one of the possible zigzagging codings of message/carrier frequency. For future work, it is suggested to explore ASSR to other possible zigzagging permutations of message/carrier frequency, and to make a comparison between the responses to stimuli with different coding types (ascending, descending and zigzagging) and frequency effects. Furthermore, this study used sinusoidal amplitude modulated sequences only. Designing and testing auditory stimuli constructed with other modulations (e.g., frequency modulation (FM), pulse width modulation (PWM), etc.) would be valuable.

In the current study, each stimulus was presented separately. Nevertheless, the stimuli in most of the compared studies were played simultaneously, which may decrease their classification performance. In other words, for each subject of the current study, after each stimulus was played, the fatigue reported by the subject was written down, then another stimulus was presented, and so on. There were two reasons for this. We wanted to: 1) ensure that whether the stimuli in each introduced set evoked adequate inherently distinguishable responses in the brain, 2) measure the amount of fatigue that each stimulus caused to each subject, so we had to present the stimuli separately (i.e., one by one). Although simultaneous presentation of the stimuli is required in BCI paradigms, this was not the case in our study, which is not a BCI paradigm. This study is a preliminary step that investigated the feasibility of utilizing the proposed stimuli in aBCI paradigm. For this purpose, the amount of inherent distinguishability between the responses in each set, along with subjects’ fatigue were measured through the separate presentation. However, in simultaneous presentation of the stimuli, f_c_ coding in the rhythmic set will help the users to focus on and discriminate between the stimuli. This implies that the classification performance of the responses in the simultaneous stimuli presentation might not be too different from our results that are obtained via separate stimuli presentation. Obviously, however, there is still a need for future works to investigate the simultaneous presentation of the proposed stimuli to see how and to what extent the results may differ from this study. More specifically, speed can only be evaluated in simultaneous presentation of the stimuli.

Each subject’s fatigue measurement was only subjective (i.e., VAS). However, for the future works, objective fatigue measurement through EEG analysis is worth conducting. Moreover, measuring each subject’s acceptance and interest rates might give more insight into their preference, and therefore stimuli development.

Since this paper is a preliminary step towards developing a novel aBCI, the sample consisted solely of healthy humans. Thus, conducting the experiment performed in this paper on completely locked-in state syndrome (CLIS) patients is proposed for future work to see whether they are useful for those individuals. Additionally, the protocol can be tested on a larger sample size.

In this study, we aimed at exploring the responses in common domains (e.g., time and frequency). However, nonlinear and/or time-frequency analyses can be performed and compared in future studies. Our protocol used 19 channels, which is not sufficient for the responses to be used in source localization algorithm. High-density recording, therefore, is recommended so that source localization would also be possible.

## Conclusion

In this study, six sinusoidal amplitude-modulated auditory stimuli with multiple message frequency coding have been introduced. Our findings suggest that both simple and rhythmic stimuli sets evoked highly distinguishable responses. Moreover, listening to the rhythmic stimuli set significantly reduced the subjects’ fatigue. Thus, testing these novel stimuli in a BCI experiment seems to be worthwhile for enhancing the number of commands by means of the multiple message frequency coding and for reducing the subjects’ fatigue.

## Acknowledgements

This work was supported by a grant from the deputy of the research review board and the ethics community of Tehran University of Medical Sciences (Grant NO: 30863)

